# Intrinsic functional connections determine how curiosity and prediction errors enhance memory

**DOI:** 10.1101/2023.04.21.537775

**Authors:** Kathrin C. J. Eschmann, Ashvanti Valji, Kim S. Graham, Andrew D. Lawrence, Matthias J. Gruber

## Abstract

Individuals differ in the way they seek information, acquire knowledge, and form memories. Neural fingerprints of intrinsic functional connectivity distinguish between individuals and predict inter-individual differences in task performance. Both curiosity – the desire to acquire new information – and information prediction errors (IPEs) – the mismatch between information and previous expectations – enhance memory but differ considerably between individuals. The present study assessed whether inter-individual differences in functional connectivity measured using resting-state fMRI determine the extent to which individuals benefit from memory-enhancing effects of curiosity and IPEs. We found a double dissociation between individual differences in mesolimbic functional connectivity, which accounted for curiosity-driven but not IPE-related memory enhancements, and individual differences in cingulo-hippocampal functional connectivity, which predicted IPE-driven but not curiosity-related memory enhancements. These novel findings on how inter-individual differences in dissociable intrinsic functional networks determine memory enhancements stress the need to account for these differences in theoretical frameworks of curiosity and memory.

## INTRODUCTION

Individuals differ in their motivation to seek, explore, and consequently remember information. Over recent years, research has shown that these individual differences are determined by individually unique neural patterns of functional connectivity measured at rest. These neural fingerprints of intrinsic functional connectivity were shown to distinguish between individuals (Finn et al., 2015), be stable over time (Horien et al., 2019; Jalbrzikowski et al., 2020; Laumann et al., 2017; Shehzad et al., 2009), and even predict unrelated task performance (Baldassarre et al., 2012; Gerraty et al., 2014). Notably, functional connectivity at rest was shown to be highly comparable to task-driven coactivation patterns (Cole et al., 2014; Gratton et al., 2016, 2018; Mennes et al., 2010; Tavor et al., 2016), indicating that these intrinsic networks support task-related cognitive processes (Cole et al., 2014; Gratton et al., 2018). For example, within the research field of learning and memory, it has been shown that intrinsic functional connections between the prefrontal cortex (PFC) and the hippocampus predict an individual’s learning behaviour (Gerraty et al., 2014). These cortico-hippocampal functional connections have been interpreted to reflect memory control processes that are necessary to encode and retrieve episodic memories (see Anderson et al., 2016; Kim, 2011; Ranganath, 2010; Simons & Spiers, 2003 for reviews). Importantly, how much information an individual seeks and remembers is also strongly driven by states of incentive salience and associated dopaminergic activity within the cortico-mesolimbic circuit (see Miendlarzewska et al., 2016; Shohamy & Adcock, 2010 for reviews). For instance, individual differences in curiosity and intrinsic mesolimbic functional connectivity were shown to drive real-life information seeking (Eschmann et al., 2023; Lydon-Staley et al., 2021). In addition, individual differences in intrinsic hippocampal-striatal functional connectivity have been linked to affect-related diversity in physical exploration (Heller et al., 2020). Similar to extrinsic rewards, high states of curiosity and underlying mesolimbic functional connectivity increase the likelihood of encountered information to be remembered later on (Duan et al., 2020; Fandakova & Gruber, 2021; Galli et al., 2018; Gruber et al., 2014; Kang et al., 2009; Lang et al., 2022; Ligneul et al., 2018; McGillivray et al., 2015; Murphy et al., 2021; Stare et al., 2018). Several states of incentive salience have been associated with activity in the cortico-mesolimbic dopaminergic circuit (e.g., Bromberg-Martin & Hikosaka, 2009; Frank et al., 2019; Gruber et al., 2014) but it remains an open question whether dissociable intrinsic functional networks reveal the degree to which an individual benefits from different states of incentive salience.

A conceptual model that summarises and integrates cognitive processes and neural underpinnings of salience-driven memory enhancement is the recently proposed Prediction, Appraisal, Curiosity, and Exploration (PACE) framework (Gruber & Ranganath, 2019). Curiosity as the intrinsic desire to acquire new information is suggested to make the associated information particularly salient in absence of any extrinsic reinforcers, thereby guiding subsequent exploration and information-seeking behaviour in order to reduce uncertainty and information gaps (Berlyne, 1960; Gottlieb et al., 2013; Gottlieb & Oudeyer, 2018; Kidd & Hayden, 2015; Loewenstein, 1994; van Lieshout et al., 2020). According to the PACE framework, curiosity leads to enhanced hippocampus-dependent encoding and consolidation of acquired information via neuromodulation of the mesolimbic dopaminergic circuit (Gruber & Ranganath, 2019). Consistent with the PACE framework, the fledgling field of curiosity research has shown that task-related curiosity states elicit activation and functional connectivity of two key regions of the mesolimbic system, namely, the ventral tegmental area (VTA) and the nucleus accumbens (NAcc), which predict curiosity-driven memory enhancements (Gruber et al., 2014; Kang et al., 2009; Lau et al., 2020; Ligneul et al., 2018; Oosterwijk et al., 2020; Poh et al., 2022). Furthermore, it has been demonstrated that large inter-individual differences in the learning- and memory-enhancing effect of curiosity exist (Fandakova & Gruber, 2021; Gruber et al., 2014; Wade & Kidd, 2019) but the functional underpinnings of these individual differences are unclear. The memory-enhancing and neuromodulatory effects of curiosity are highly similar to the effects of extrinsic rewards on memory. Research on reward further suggests that individual differences in reward-related memory enhancements are determined by VTA-NAcc and VTA-hippocampal structural connectivity (Elliott et al., 2022; Reggente et al., 2018) as well as functional connectivity within the cortico-mesolimbic dopaminergic system irrespective of whether it is measured during a task or at rest (Frank et al., 2019). Based on these findings and following the rationale that intrinsic functional connections measured at rest support cognitive performance due to overlap with task-related coactivation patterns (Cole et al., 2014; Gratton et al., 2018), individual differences in the strength of intrinsic functional connectivity within the mesolimbic system might predict the magnitude of curiosity-driven memory enhancements.

Recent research has shown that it is not only curiosity about information but also the discrepancy between the actual information and the expectancy thereof, measured as an information prediction error (IPE), that affects encoding and later memory (Marvin & Shohamy, 2016). Information that is perceived as more interesting or rewarding than previously expected is associated with a positive IPE and remembered better than information that is less interesting than expected and hence linked to a negative IPE. Behavioural studies have shown that positive IPEs enhance memory independently of curiosity (Fandakova & Gruber, 2021; Fastrich et al., 2018; Lang et al., 2022; Marvin & Shohamy, 2016) and that individual differences in IPE-driven memory enhancements start to emerge during adolescence (Fandakova & Gruber, 2021). However, the underlying functional networks of IPE -driven memory enhancements are not known. Consistent with research on reward prediction errors, it has been proposed that IPEs – similarly to curiosity – also depend on the mesolimbic dopaminergic circuit (Bromberg-Martin & Hikosaka, 2009, 2011; Brydevall et al., 2018; Charpentier et al., 2018; van Lieshout et al., 2018), suggesting that novel information itself can act as a reward signal (Barto et al., 2013; Marvin & Shohamy, 2016; van Lieshout et al., 2020). This idea is supported by electrophysiological studies in non-human primates showing increased firing of midbrain dopamine and lateral habenula neurons for both information and reward prediction errors (Bromberg-Martin & Hikosaka, 2009, 2011; Kobayashi & Hsu, 2019; Matsumoto & Hikosaka, 2007). Similarly, initial electrophysiological evidence in humans suggests that IPEs are processed in reward circuitries by revealing that both information and reward prediction errors independently alter the feedback-related negativity (Brydevall et al., 2018). Despite this evidence, a direct link between neuromodulation of the mesolimbic dopaminergic circuit and IPE-driven memory enhancements is still missing.

Alternatively and in accordance with the PACE framework (Gruber & Ranganath, 2019), it has been proposed that discrepancies between actual and expected outcome can be detected by both the hippocampus and the anterior cingulate cortex (ACC). Regarding the role of the hippocampus, theoretical frameworks have suggested that the hippocampus forms cognitive maps that help to generate context-based predictions and code violations thereof (O’Keefe & Nadel, 1978). Furthermore, it has been theorised that the hippocampus also enables memory-based predictive coding of non-spatial information (Barron et al., 2020; Stachenfeld et al., 2017). Indeed, activation of the hippocampus has been shown to increase in response to the detection of prediction errors (Chen et al., 2011; Dickerson et al., 2011; Duncan et al., 2009, 2012; Kumaran & Maguire, 2006; Schiffer et al., 2012; Vinogradova, 2001). Notably, latest research has demonstrated that prediction errors coded in the hippocampus enable learning and memory updating by shifting hippocampal states from processing of ongoing predictions to information intake, allowing memory encoding based on prediction errors to take place (Bein et al., 2020; Sinclair et al., 2021). With regard to the role of the ACC, numerous theoretical frameworks have pointed out that the ACC is responsible for conflict monitoring (Botvinick, 2007; Botvinick et al., 2001) and outcome prediction (Alexander & Brown, 2011; Brown & Braver, 2005; Monosov et al., 2020; Vassena et al., 2020). Specifically, ACC activity has been suggested to signal the degree to which information is unexpected or surprising, leading to adjustments in cognitive control that resolve the detected conflict and to updated predictions for future encounters (Alexander & Brown, 2011; Vassena et al., 2020). Accumulating evidence supports this idea by showing greater ACC activation for unexpected information irrespective of valence (Braver et al., 2001; Ferdinand & Opitz, 2014; Wessel et al., 2012). Taken together, it remains to be investigated whether intrinsic functional connectivity within the mesolimbic dopaminergic circuit or between the hippocampus and ACC drives IPE-related memory enhancements.

The present study aimed at disentangling the functional connections that determine individual differences in curiosity-related and IPE-driven memory enhancements. A final sample of 80 participants took part in a trivia paradigm along with a separate 10-minute resting-state functional magnetic resonance imaging (fMRI) scan (Figure 1). During the trivia paradigm, participants were presented with a series of trivia questions and the associated answers, to which they had previously rated their curiosity. After presentation of the trivia question, participants were asked to indicate how interesting they found the answer and IPEs were calculated as the difference between the interest and initial curiosity rating for each trivia answer (interest rating – curiosity rating; Marvin & Shohamy, 2016), resulting in positive, no, or negative IPEs. Memory of the trivia answers was tested in a surprise cued recall 24 hours later. Based on previous research, we expected trivia answers associated with high curiosity (rating 4-6) to be remembered better than trivia answers linked to low curiosity (rating 1-3). Independently, trivia answers associated with positive IPEs were expected to be remembered better than trivia answers related to no or negative IPEs. In order to link intrinsic functional connectivity to behavioural performance, memory enhancements were calculated as differences in recall for the conditions of curiosity (high curiosity – low curiosity) and IPEs (positive IPEs – negative IPEs).

**Figure 1.**
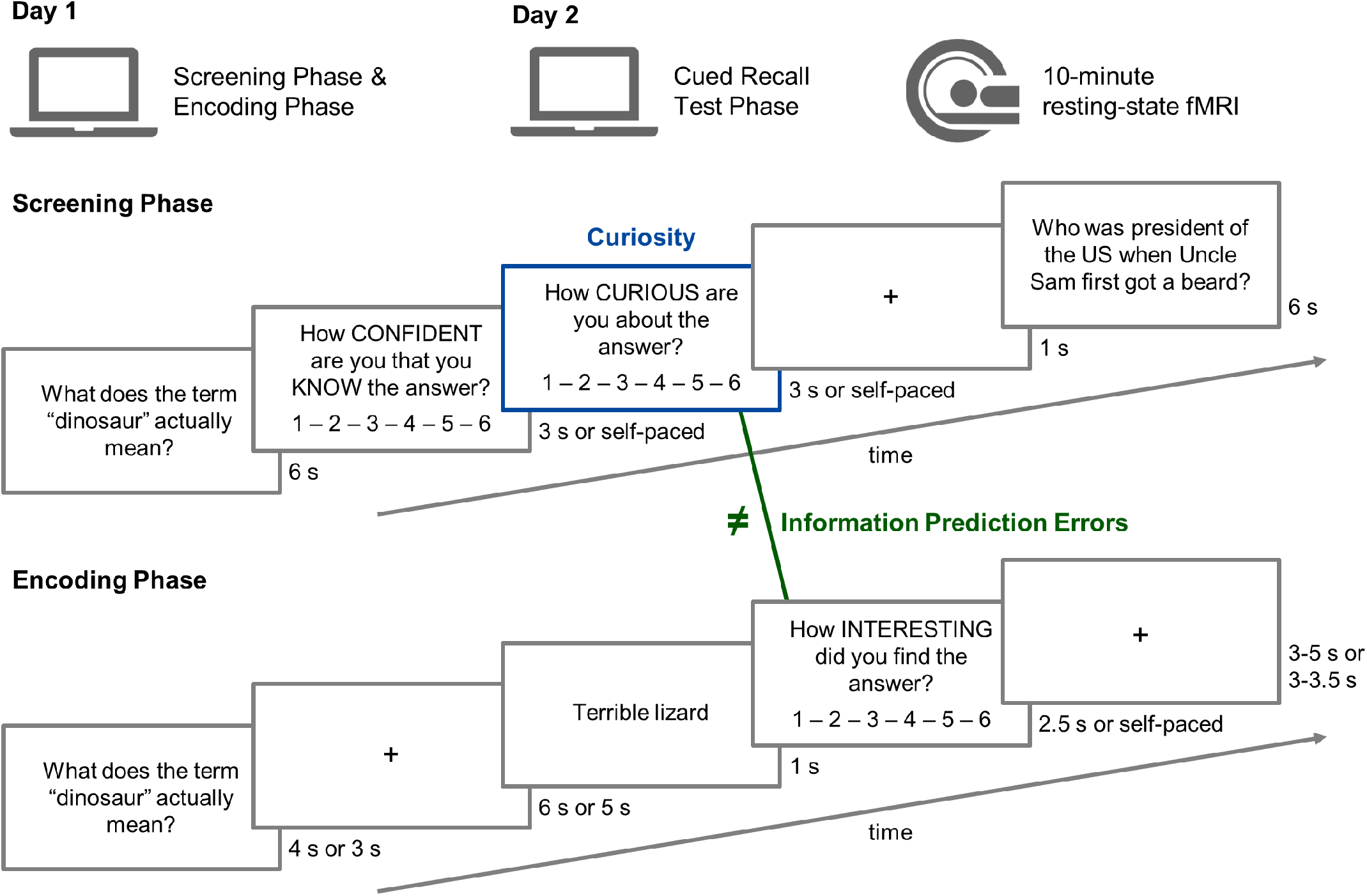
Overview of the trivia paradigm and the separate resting-state fMRI scan. During the first day, participants performed the screening and encoding phase of the trivia paradigm. A day later, participants’ memory performance was assessed in a cued recall test phase. Either after the memory test or on a separate day, a 10-minute resting-state fMRI scan was conducted. During the screening phase, participants rated their knowledge and curiosity about trivia answers after seeing the associated trivia questions. The screening phase continued until the same target amount of low curiosity (rating 1-3) and high curiosity (rating 4-6) trivia answers was reached. During the encoding phase, participants saw the trivia question, anticipated the respective trivia answer during a delay period, were presented with the trivia answer, and were asked to rate how interesting they found the presented trivia answer. Positive and negative information prediction errors (IPEs) occurred when the interest rating was higher or lower than the curiosity rating about the same trivia answer in the screening phase, respectively. Different presentation times in the trivia paradigm are because participants of two neuroimaging studies were combined for the present investigation (*n* = 55 and *n* = 40). Within the final sample of 80 participants, no memory performance differences were found between participants of the two neuroimaging studies.

For analyses of intrinsic functional connectivity, regions of interest (ROIs) were selected based on previous research summarised in the PACE framework (Gruber & Ranganath, 2019). Consequently, ROI-to-ROI functional connectivity values between left/right VTA, left/right NAcc, bilateral ACC, left/right hippocampus, and left/right lateral prefrontal cortex (LPFC) were calculated for each participant, whereby each ROI served as a seed ROI once and otherwise as a target ROI (Figure 2). In order to correct for multiple comparisons, the false discovery rate (FDR) seed-level correction was applied and adjusted *p*-values are reported (Benjamini & Hochberg, 1995). We hypothesised that individual differences in intrinsic functional connectivity between VTA and NAcc should predict curiosity-driven memory enhancements. If it holds true that IPEs resemble reward prediction errors, it seems likely that VTA-NAcc intrinsic functional connectivity also relates to IPE-driven memory enhancements. However, if IPE-related memory enhancements are driven by IPEs processed in the ACC and hippocampus, it might be possible that individual differences in intrinsic functional connectivity between the ACC and hippocampus predict IPE-driven memory enhancements.

**Figure 2.**
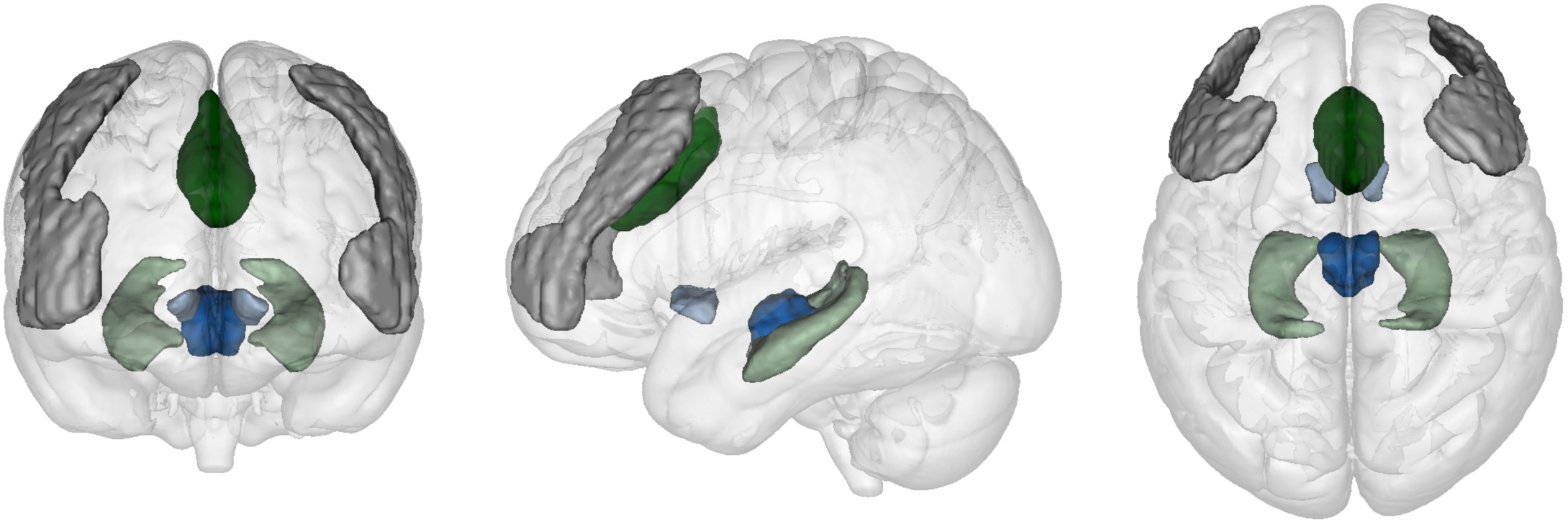
Coronal, sagittal, and axial view of the ROIs from the PACE framework that were used for intrinsic functional connectivity analysis. The ventral tegmental area (VTA) is displayed in dark blue, the nucleus accumbens (NAcc) in bright blue, the anterior cingulate cortex (ACC) in dark green, the hippocampus in bright green, and the lateral prefrontal cortex (LPFC) in grey. The figure was created with the multi-image analysis GUI Mango (ric.uthscsa.edu/mango).

## MATERIALS AND METHODS

### Participants

Overall, 95 healthy participants took part in two neuroimaging studies that were combined for the present investigation (*n* = 55 and *n* = 40). All participants were recruited amongst Cardiff University’s student population. Data of 15 participants were excluded from statistical analyses due to the following issues: Ten participants showed excess motion artefacts during fMRI data acquisition, for four participants behavioural data of the trivia paradigm were missing due to technical issues, and one participant demonstrated poor memory performance (< 10% overall memory accuracy). The final sample consisted of 80 right-handed participants (63 female, 17 male, *M*age = 19.8 years, age range = 18-29 years) with normal or corrected-to-normal vision. Notably, overall memory performance did not differ between participants of the two neuroimaging studies (*t*(78) = 0.20, *p* = .841, Cohen’s *d* = 0.05, two-tailed). Furthermore, when including the factor study as a control variable in separate one-way ANCOVAs, the reported memory-enhancing effects of curiosity (*F*(1,157) = 51.49, *p* < .001, η^2^ = .25) and IPEs (*F* (2,236) = 13.69, *p* < .001, η^2^ = .10) remained significant. Participants gave written informed consent prior to participation and received either course credits and/or payment as reimbursement. Ethics were approved by the Cardiff University School of Psychology Research Ethics Committee in accordance with the declaration of Helsinki.

### Experimental design

All participants took part in a trivia paradigm and a separate 10-minute resting-state fMRI scan. The trivia paradigm consisted of a screening, an encoding, and a memory recall phase (Figure 1). During the screening phase, trivia questions were randomly selected from a pool of trivia questions and presented for six seconds. Afterwards, participants were asked to indicate on a 6-point Likert scale how likely it was that they *know* the respective trivia answer (extremes of scale: 1 = “not at all confident” and 6 = “extremely confident”). Questions, to which participants were confident of knowing the answer (rating of 6), were discarded from the following encoding phase, ensuring that the effects of curiosity and IPEs on memory were not confounded by previous knowledge. Participants were also asked to rate on a 6-point Likert scale how *curious* they were about the trivia answer (extremes of scale: 1 = “not at all curious” and 6 = “extremely curious”). Trivia questions with responses of 1-3 were assigned to the pool of low curiosity trials whereas trivia questions with a rating of 4-6 were allocated to the pool of high curiosity trials. To remove an initial bias in the curiosity rating, the first four trials of the screening phase were not included in the pool of trials for the encoding phase irrespective of the given rating responses. Participants were asked to rest their fingers on the relevant keys on the keyboard and encouraged to use the full range of the scales for both the knowledge and curiosity ratings. The screening phase terminated when the same target number of trivia questions was assigned to each pool of high and low curiosity trials.

During the subsequent encoding phase, participants were presented with an equal number of high and low curiosity trivia questions and their respective answers. The presentation of a trivia question was followed by a fixation cross, during which participants were asked to anticipate the upcoming answer. Subsequently, the respective trivia answer was presented for one second followed by an interest rating that asked participants how *interesting* they found the answer on a 6 -point Likert scale (extremes of scale: 1 = “not at all interesting” and 6 = “extremely interesting”). During the subsequent inter-trial interval, a fixation cross was presented. The encoding phase consisted of four blocks separated by three self-paced breaks and lasted approximately 30-40 minutes.

Roughly 24 hours after the encoding phase, participants returned to the laboratory to complete a surprise cued recall memory test. Trivia questions from the encoding phase were presented in a random order and participants were encouraged to remember and type in the respective trivia answer as accurately as possible. The memory recall phase took participants approximately 20-40 minutes to complete.

Even though the overall design of the trivia paradigm was the same for both neuroimaging studies that were combined for the present investigation, some characteristics differed. In the screening phase, questions were drawn from a pool of either 294 or 425 trivia questions and either 60 or 140 trials were selected based on the curiosity ratings for the encoding phase. The time, during which the knowledge and curiosity ratings were presented and in which participants were able to give their responses, was three seconds in the *n* = 55 study and self-paced in the *n* = 40 study. In the encoding phase of the *n* = 55 study, 30 high and 30 low curiosity trivia questions were presented for four seconds in alternating sets of five consecutive high or low curiosity trials, whereby the set order was counterbalanced across participants. In the *n* = 40 study, 70 high and 70 low curiosity trivia questions were presented intermixed for three seconds each. The fixation cross that followed the trivia question was presented for either five or six seconds. For the interest rating, participants had either 2.5 seconds to give their response or the responses were self-paced. The inter-trial interval consisted of a jittered time interval of either 3-5 or 3-3.5 seconds. The *n* = 55 study, additionally consisted of the presentation of a one-second exclamation mark, a two-second image of a face, and a four-second fixation cross that preceded the presentation of each trivia question (not shown in Figure 1). Given that face images were not presented in the *n* = 40 study, memory of incidentally presented faces was not investigated.

### fMRI acquisition

Imaging data were obtained at CUBRIC, Cardiff University, using a Siemens Magnetom Prisma 3T MRI scanner with a 32-channel head coil. High-resolution T1-weighted structural images were obtained using an MPRAGE sequence (TR = 2500 ms, TE = 3.06 ms, flip angle = 9°, FoV = 256 mm^2^, voxel-size = 1 mm^3^, slice thickness = 1 mm, 224 sagittal slices, bandwidth = 230 Hz/pixel; acquisition time = 7.36 min). During the structural scan, participants watched a film to help reduce movement, boredom, and nervousness. For resting-state fMRI, 50 transversal slices were acquired by using an echoplanar imaging (EPI) sequence (TR = 3000 ms, TE = 30 ms, flip angle 89°, FoV = 192 mm^2^, voxel-size = 2 mm^3^, slice thickness = 2 mm, bandwidth = 2170 Hz/pixel; acquisition time = 10.11 min). A black fixation cross centred on a grey background was presented during resting-state fMRI acquisition. Participants were instructed to keep their eyes open, fixate on the cross, and try to the best of their ability to keep their minds clear.

### Resting-state functional connectivity preprocessing and analysis

Resting-state fMRI data were pre-processed using the CONN toolbox (version 18b; Whitfield-Gabrieli and Nieto-Castanon, 2012), in conjunction with the Statistical Parametric Mapping (SPM12) modules (Wellcome Trust Centre for Neuroimaging, London) executed in MATLAB (version 2015). In a first step, functional scans were realigned and resampled to a reference structural image using SPM12 (Andersson *et al*., 2001) and slice-time corrected (Henson *et al*., 1999), adjusting for differences in acquisition times between the inter-leaved scans. The Artefact Detection Tool (ART) was used to flag potential outliers with a framewise displacement > 0.5 mm and global BOLD signal change exceeding three standard deviations of subject-specific means. Then, structural and functional images were normalised into MNI space and segmented (Ashburner and Friston, 2005), with 2 mm isotropic voxels for functional and 1 mm isotropic voxels for structural images. Functional imaging was spatially smoothed using 6 mm full-width-half maximum (FWHM) Gaussian Kernel. Next, images were denoised using CONN’s anatomical component noise correction procedure (Behzadi *et al*., 2007; Chai *et al*., 2012). Twelve noise components (three translation, three rotation parameters, and their respective first-order derivatives) were identified (Friston *et al*., 1996) to reduce motion variability in the BOLD signal. Outlier scans identified in ART were scrubbed during this step. To remove slow trends in the signal and initial magnetization transients from the BOLD signal, a linear detrending was used. To keep as much data of interest as possible, the default settings in the CONN toolbox were changed so that global signal change was not removed. Finally, the standard band-pass filter between 0.008 Hz and 0.09 Hz was used. In addition, data from participants with over 15% of invalid scans as identified by ART were removed from analysis. According to this exclusion criterion, data from ten participants were removed.

To calculate intrinsic functional connectivity between the brain regions of the PACE framework, masks of left/right VTA, left/right NAcc, bilateral ACC, left/right hippocampus, and left/right LPFC were used as regions of interest (ROIs; Figure 2). The bilateral mask of the VTA was taken from Murty *et al*. (2014) and separated into left and right VTA masks in FSL (Jenkinson *et al*., 2012). Masks of left and right NAcc and left and right hippocampus were taken from the Harvard-Oxford cortical atlas integrated in the CONN toolbox. The bilateral ACC mask and left and right LPFC masks were taken from the Harvard-Oxford networks atlas in CONN. The BOLD time series of each ROI was computed by averaging the voxel time series across all voxels within the ROI. Each participant’s intrinsic functional connectivity values were computed as Fisher’s z-transformed bivariate Pearson correlation coefficients between seed and target ROI’s BOLD time series.

### Data analyses

Statistical analyses of behavioural data were carried out with base and stats functions in RStudio (RStudio Team, 2022) and effect sizes were calculated with the R package effectsize (Ben-Shachar et al., 2020). Result figures were created with the R packages ggplot2 (Wickham, 2009) and superb (Cousineau et al., 2021). The effect of curiosity on later memory was investigated with a one-sample *t*-test comparing memory accuracy for trivia answers associated with levels of high and low curiosity, respectively. The effect of IPEs on memory accuracy was explored with a one-way repeated-measures ANOVA with the within-subject factor IPEs (positive vs. no vs. negative). Memory accuracy was determined as the percentage of recalled trivia answers for each level of the two conditions. IPEs were calculated as the difference between the interest and curiosity rating for each trivia answer (interest rating – curiosity rating; Marvin & Shohamy, 2016). Positive IPEs were defined by greater interest in the trivia answer than previously indicated in the curiosity rating whereas negative IPEs were revealed by lower interest in the trivia answer than previously indicated in the curiosity rating. For the calculation of IPEs, trials, for which the curiosity and interest rating did not differ, were specified as no IPE trials. Furthermore, trials, for which no interest rating was given or with a response time < 50 ms for the interest rating, were classified as no IPE trials. Effects of curiosity and IPEs on later memory were investigated separately because more positive IPEs were present in low curiosity trials whereas more negative IPEs appeared in high curiosity trials, leading to non-normally distributed data. Post hoc *t*-tests were calculated to further explore a significant main effect.

The positive relationship of intrinsic functional connectivity with curiosity-driven and IPE-driven memory enhancements was assessed with separate one-tailed linear regression analyses conducted in the CONN toolbox (version 18b; Whitfield-Gabrieli & Nieto-Castanon, 2012), in conjunction with the Statistical Parametric Mapping (SPM12) modules (Wellcome Trust Centre for Neuroimaging, London) executed in MATLAB (version 2015). Memory enhancements were calculated as memory accuracy differences for the conditions of curiosity (high curiosity – low curiosity) and IPEs (positive IPEs – negative IPEs). Functional connectivity values of all ROI-to-ROI connections were entered as predictors into regression analyses with the curiosity-driven memory enhancement as dependent variable. The same analyses were conducted with the IPE-driven memory enhancement as the dependent variable. For all analyses, the significance level was set to α = .05 and if not indicated differently, one-tailed results are reported due to the directional hypotheses. For effect sizes of between- and within-subject *t*-tests, Cohen’s *d* and *dz* values are reported, respectively. In order to correct for multiple comparisons, the false discovery rate (FDR) method was applied and adjusted *p*-values were reported (Benjamini & Hochberg, 1995). For analyses of the relationship between ROI-to-ROI functional connectivity and memory enhancements, an FDR seed-level correction threshold was used. To avoid biases from outliers for all statistical analyses, outliers were detected with the Tukey’s method using three interquartile ranges (Tukey, 1977). No data had to be excluded based on this outlier detection.

## RESULTS

### Curiosity and IPEs enhance memory independently

As expected, participants remembered trivia answers associated with high curiosity ratings better than trivia answers associated with low curiosity ratings (*t*(79) = 11.01, *p* < .001, Cohen’s *dz* = 1.23; Figure 3A). Curiosity enhanced memory accuracy by *M* = 15.29%, *SD* = 12.42%. Additionally, IPEs also enhanced memory accuracy of trivia answers as indicated by a significant main effect of IPEs (*F*(2,237) = 13.72, *p* < .001, η^2^ = .10; Figure 3B). Post hoc *t*-tests revealed that trivia answers associated with positive IPEs, that is, greater interest after presentation of the trivia answer than previously indicated in the curiosity rating, were better recalled than trivia answers linked to negative IPEs, that is, less interest than previously reported (*t*(79) = 6.45, *p* < .001, Cohen’s *dz* = 0.72). The IPE-driven memory enhancement added up to *M* = 12.24%, *SD* = 16.97%. Trivia answers associated with positive IPEs were also better remembered by *M* = 9.59%, *SD* = 17.07%, than trivia answers that were linked to no IPEs (*t*(79) = 5.03, *p* < .001, Cohen’s *dz* = 0.56). There was no difference in memory accuracy between trivia answers associated with no IPEs and negative IPEs (*t*(79) = 1.44, *p* = .077, Cohen’s *dz* = 0.16).

**Figure 3.**
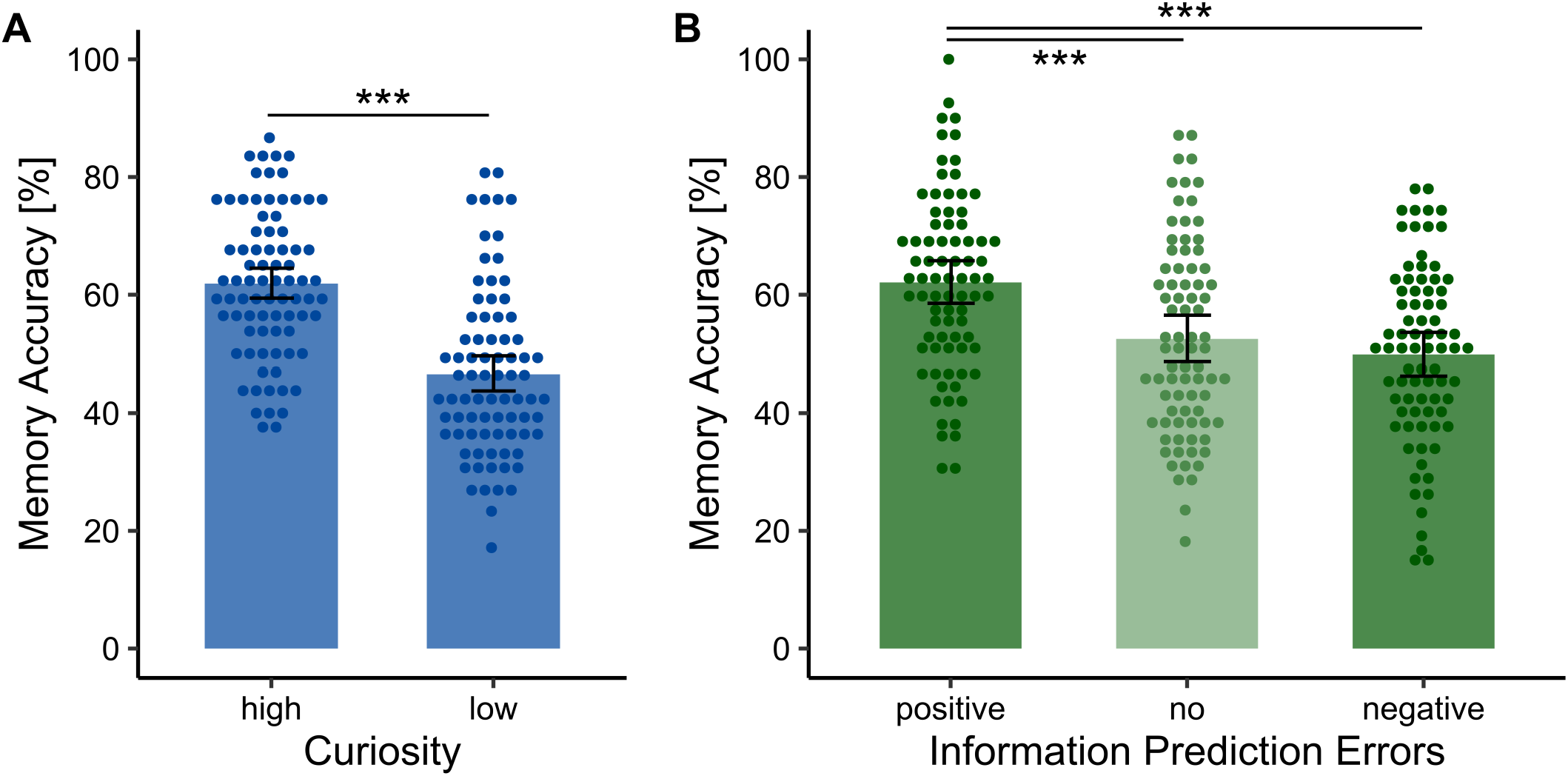
Curiosity and IPEs enhance memory. (A) Memory accuracy of trivia answers associated with high curiosity was better compared to memory accuracy of trivia answers associated with low curiosity. (B) Positive IPEs lead to higher memory accuracy of trivia answers compared to negative IPEs. Trivia answers associated with positive IPEs were also better remembered than trivia answers associated with no IPEs. There was no difference in memory accuracy between no IPEs and negative IPEs. Error bars indicate correlation- and difference-adjusted 95% confidence intervals of the mean for within-subject designs (Cousineau et al., 2021). * *p* < .05, ** *p* < .01, *** *p* < .001

### Individual differences in mesolimbic and cingulo-hippocampal functional connectivity dissociate between curiosity-driven and IPE-related memory enhancements

To investigate the positive relationship of cortico-mesolimbic functional connectivity with memory enhancements, one-tailed seed-level corrected linear regressions with intrinsic functional connectivity between ROIs were calculated separately for curiosity-driven and IPE-driven memory enhancements, respectively. Thereby, a double dissociation between mesolimbic and cingulo-hippocampal intrinsic functional connectivity was found. For curiosity-driven memory enhancements, individual differences in functional connectivity between left VTA and right NAcc predicted the magnitude of curiosity-driven memory enhancements (β = 0.33, *t* (78) = 2.65, FDR-adjusted *p* = .039; Figure 4A). Similarly, intrinsic functional connectivity between right NAcc and right VTA was associated with curiosity-driven memory enhancements (β = 0.25, *t* (78) = 2.39, FDR-adjusted *p* = .038; Figure 4B-4C). In contrast, both left VTA – right NAcc and right NAcc – right VTA functional connectivity was not associated with IPE-driven memory enhancements (FDR-adjusted *p*-values > .744; Figure 4D-4F). None of the other functional connections predicted curiosity-driven memory enhancements (FDR-adjusted *p*-values > .146).

**Figure 4.**
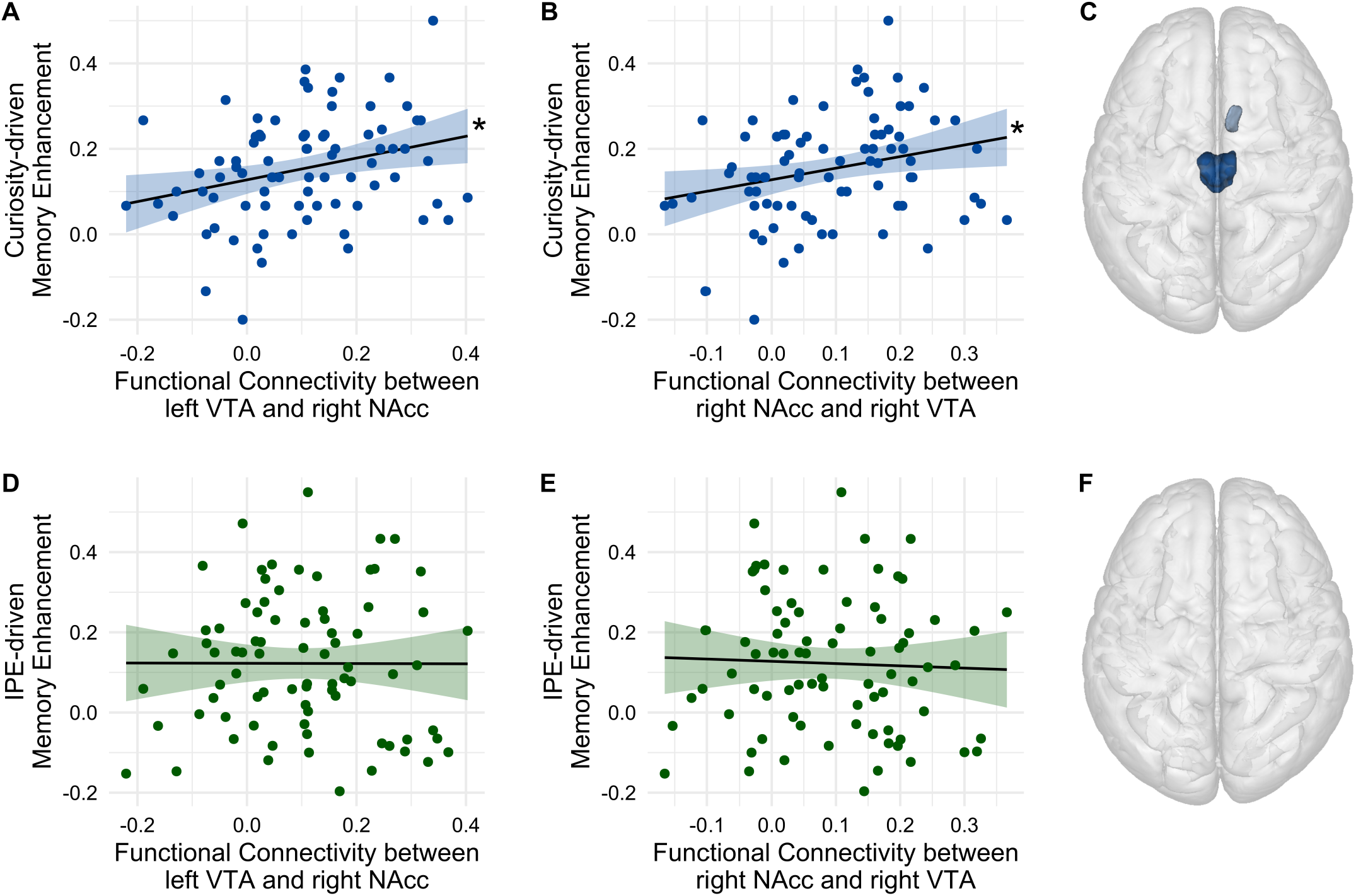
Relationship of intrinsic mesolimbic functional connectivity with curiosity-driven and IPE-driven memory enhancements. (A) Individual differences in intrinsic left VTA – right NAcc functional connectivity predicted the magnitude of curiosity-driven memory enhancements. (B) Right NAcc – right VTA functional connectivity was positively associated with curiosity-related memory enhancement. (C) Axial view of the mesolimbic ROIs that showed functional connectivity related to curiosity-driven memory enhancements, namely left/right VTA (dark blue) and right NAcc (bright blue). (D) Individual differences in left VTA – right NAcc functional connectivity did not predict IPE-driven memory enhancements. (E) Right NAcc – right VTA functional connectivity was not associated with IPE-related memory enhancement. (F) No mesolimbic ROIs showed functional connectivity associated with IPE-driven memory enhancements. * *p* < .05, ** *p* < .01, *** *p* < .001

For IPE-driven memory enhancements, individual differences in functional connectivity between bilateral ACC and left hippocampus predicted the magnitude of IPE-driven memory enhancements (β = 0.31, *t*(78) = 3.40, FDR-adjusted *p* = .004; Figure 5D). Furthermore, intrinsic functional connectivity between bilateral ACC and right hippocampus was associated with IPE-driven memory enhancements (β = 0.28, *t*(78) = 2.82, FDR-adjusted *p* = .012; Figure 5E-5F). Contrarily, both bilateral ACC – left hippocampus and bilateral ACC – right hippocampus functional connectivity did not predict curiosity-driven memory enhancements (FDR-adjusted *p*-values > .575; Figure 5A-5C). None of the other connections predicted IPE-driven memory enhancements (FDR-adjusted *p*-values > .051). Altogether, these results suggest that individual differences in intrinsic mesolimbic and cingulo-hippocampal functional connectivity predict how curiosity and IPEs enhance memory, respectively.

**Figure 5.**
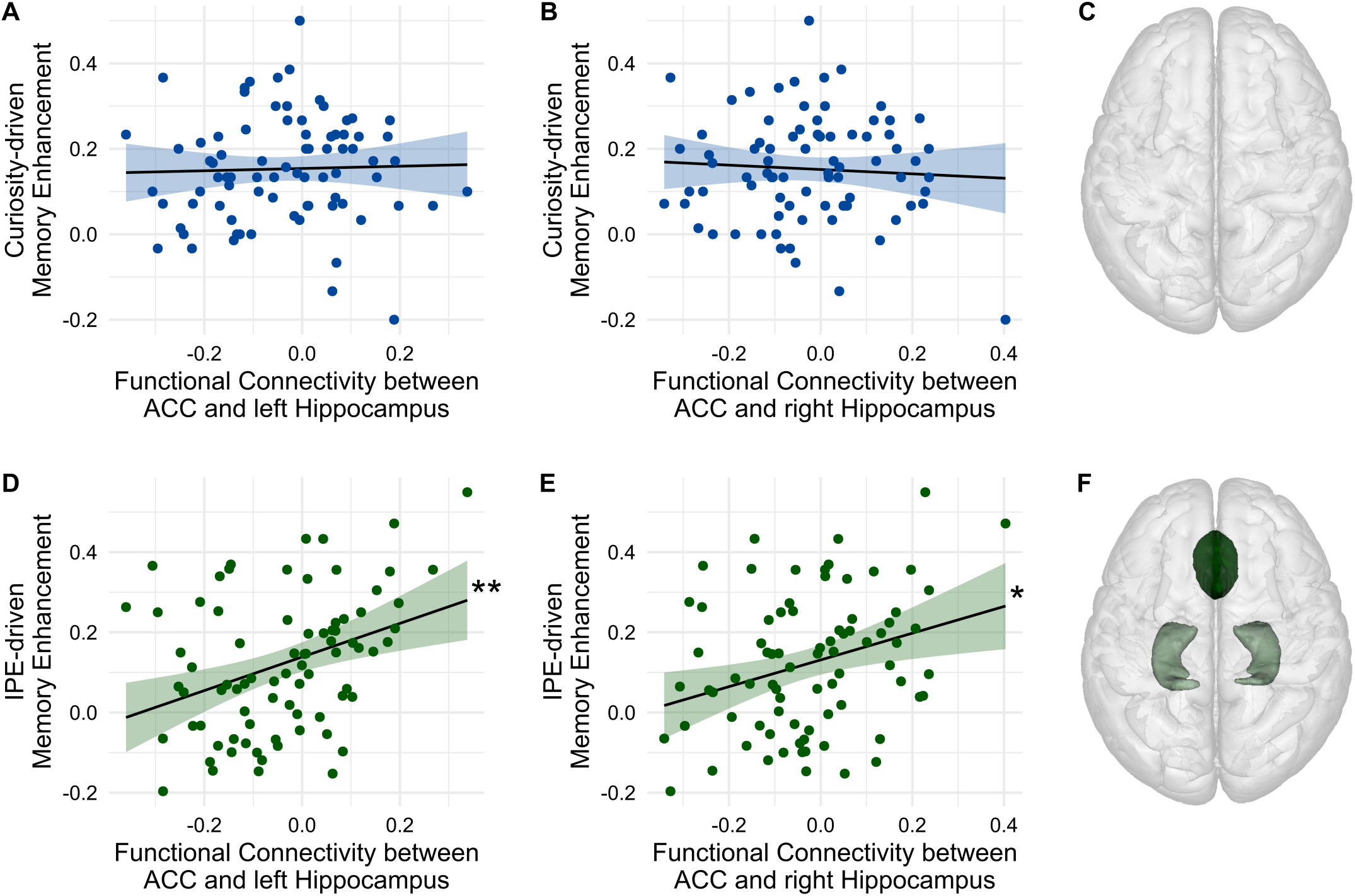
Relationship of intrinsic cingulo-hippocampal functional connectivity with curiosity-driven and IPE -driven memory enhancements. (A) Individual differences in bilateral ACC – left hippocampus functional connectivity predicted the magnitude of IPE-driven memory enhancements. (B) Bilateral ACC – right hippocampus functional connectivity was positively associated with IPE-related memory enhancement. (C) No cingulo-hippocampal ROIs showed functional connectivity related to curiosity-driven memory enhancements. (D) Individual differences in bilateral ACC – left hippocampus functional connectivity did not predict curiosity-driven memory enhancements. (E) Bilateral ACC – right hippocampus functional connectivity was not associated with curiosity-related memory enhancement. (F) Axial view of the cingulo-hippocampal ROIs that showed functional connectivity associated with IPE-driven memory enhancements, namely ACC (dark green) and left/right hippocampus (bright green). * *p* < .05, ** *p* < .01, *** *p* < .001

### Overall memory performance is associated with LPFC-hippocampal functional connectivity

To explore whether the reported findings are unrelated to overall memory performance, we investigated whether any of the functional connectivity measures predicted overall memory of the trivia answers. Therefore, one-tailed seed-level corrected linear regressions with intrinsic functional connectivity between ROIs as predictor of overall memory performance were calculated. Interestingly, individual differences in functional connectivity between left LPFC and right hippocampus predicted the magnitude of overall memory performance (β = 0.38, *t*(78) = 2.68, FDR-adjusted *p* = .036; Figure 6). This finding indicates that participants with a stronger intrinsic LPFC-hippocampal functional connectivity remembered trivia answers generally better than participants with weaker functional connectivity between LPFC and hippocampus. Importantly, neither intrinsic VTA-NAcc nor ACC-hippocampal functional connectivity predicted overall memory performance (FDR-adjusted *p*-values > .840). Furthermore, none of the other functional connections between ROIs predicted overall memory performance (FDR-adjusted *p*-values > .111). Together, these results indicate that the reported double dissociation between mesolimbic and cingulo-hippocampal functional connectivity is not driven by overall memory performance.

**Figure 6.**
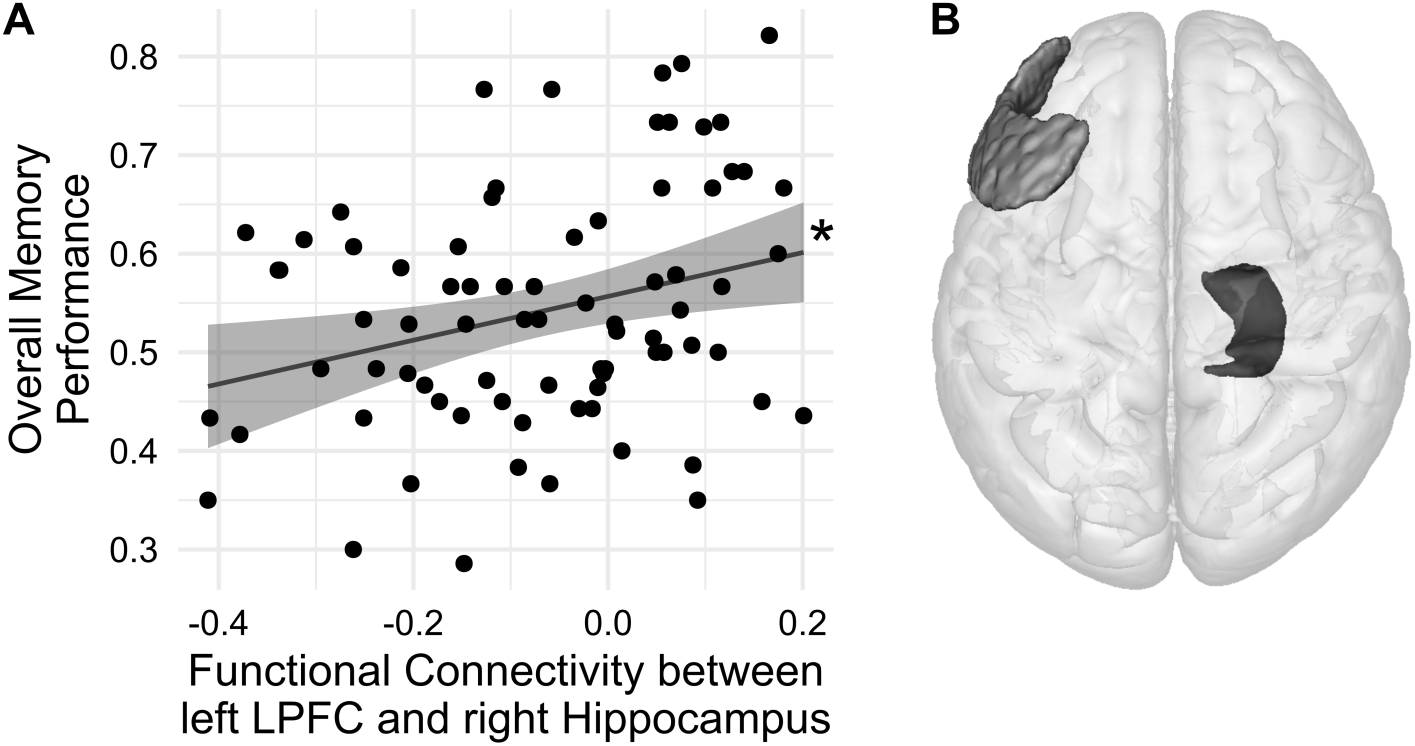
(A) Positive relationship between overall memory performance and intrinsic functional connectivity between left LPFC and right hippocampus. (B) Axial view of the ROIs that show functional connectivity associated with overall memory performance, namely left LPFC (bright grey) and right hippo-campus (dark grey). * *p* < .05, ** *p* < .01, *** *p* < .001

## DISCUSSION

The present study is the first to demonstrate a link between individual differences in the strength of task-free, intrinsic functional connectivity and the magnitude of curiosity-driven and IPE-related memory enhancements. Strikingly, mesolimbic functional connectivity was associated with curiosity-driven memory enhancements whereas cingulo-hippocampal functional connectivity was linked to IPE-driven memory enhancements, but not vice versa. This double dissociation suggests that the mechanisms by which curiosity and IPEs increase memory performance are determined by dissociable intrinsic functional networks, supporting the idea that intrinsic networks support task-related cognitive processes (Cole et al., 2014; Gratton et al., 2018). Moreover, the findings indicate that IPEs – unlike curiosity and reward prediction errors – might not rely on the mesolimbic circuit, but instead enhance memory via cingulo-hippocampal functional connectivity. Additionally, the reported findings were not driven by overall memory performance, which was predicted by LPFC-hippocampal functional connectivity, specifying the relationship of mesolimbic and cingulo-hippocampal functional connectivity with curiosity-driven and IPE-related memory enhancements, respectively. Together, these findings highlight how individual differences in intrinsic functional connectivity determine the magnitude of memory enhancements, stressing the importance of taking the effects of both curiosity and IPEs into account when investigating learning and memory.

Our findings on the relationship between VTA -NAcc functional connectivity and curiosity-related memory enhancements align with the theoretical predictions of the PACE framework and previous task-based fMRI studies on curiosity (Gruber & Ranganath, 2019). Specifically, intrinsic functional connectivity between VTA and NAcc was associated with curiosity-driven memory enhancements. This finding adds to a large variety of studies demonstrating task-related VTA and NAcc activation in response to curiosity-related learning (Gruber et al., 2014; Kang et al., 2009; Lau et al., 2020; Ligneul et al., 2018; Oosterwijk et al., 2020; Poh et al., 2022). The inter-individual variability of intrinsic mesolimbic functional connectivity in the present study may explain the individual differences that are observed in task-related mesolimbic activity and curiosity-driven memory enhancements. Research on neural fingerprints of functionally connectivity measured at rest suggests that intrinsic functional networks support task-related cognitive processes due to their high overlap with task-related coactivation patterns (Cole et al., 2014; Gratton et al., 2016, 2018; Mennes et al., 2010; Tavor et al., 2016) and their positive association with task performance (Baldassarre et al., 2012; Gerraty et al., 2014). Furthermore, functional networks have been linked to underlying brain structure based on research showing that variability in intrinsic functional connectivity is largely explained by the strength of structural pathways (see Suárez et al., 2020 for review). With regard to the mesolimbic dopaminergic circuit, tract tracing studies in rodents and monkeys have shown that the VTA and NAcc are directly and reciprocally interconnected (e.g., Avegno & Gilpin, 2022; Geisler & Zahm, 2005; Loughlin & Fallon, 1982; see Haber & Behrens, 2014 for review). In animal studies, stimulation of the VTA led to physiologically relevant dopamine release in both the ipsilateral and the contralateral ventral striatum, demonstrating the functional role of these dopamine projections (Fox et al., 2016; Ioanas et al., 2022). In line with these findings, individual differences in intrinsic functional connectivity between left VTA and right NAcc as well as right VTA and right NAcc were positively associated with individual differences in curiosity-driven memory enhancements in the present study, emphasising the importance of both ipsilateral and contralateral functional connections in the mesolimbic system for curiosity-driven learning. Microstructure indices have provided evidence for structural connectivity of the VTA and NAcc also in humans (see Jbabdi & Behrens, 2013 for review). Furthermore, the strength of both VTA-NAcc and VTA-hippocampal structural connectivity (Elliott et al., 2022; Reggente et al., 2018) and variability in cortico-mesolimbic intrinsic functional connectivity (Frank et al., 2019) have been related to individual differences in reward-related memory enhancements. The convergence of structural connectivity, intrinsic functional connectivity, and task-related measures is consistent with the idea that intrinsic functional networks are primarily determined by structural connections, providing a functionally stable basis for the support of task-related cognitive processes (Cole et al., 2014; Gratton et al., 2018). Moreover, it suggests that the individual differences in intrinsic mesolimbic functional connectivity in the present study may be based on variability in mesolimbic microstructure, causing individual differences in curiosity-driven memory enhancements. Potentially consistent with this idea, we previously found that the level of trait curiosity is associated with individual differences in fornix microstructure (Valji et al., 2019), a critical anatomical structure within the mesolimbic circuit (Shin et al., 2019).

According to the PACE framework (Gruber & Ranganath, 2019), curiosity leads to increased exploration and information seeking followed by enhanced hippocampus-dependent encoding and memory consolidation via neuromodulation of the mesolimbic circuit. While the present study shows evidence for the link between mesolimbic functional connectivity and curiosity-driven memory, we were previously able to show in a subset of participants from the present study that the strength of intrinsic VTA-NAcc functional connectivity was also positively associated with the frequency of real-life information seeking (Eschmann et al., 2023), suggesting that intrinsic mesolimbic functional connectivity primarily promotes the drive to seek new information (see FitzGibbon et al., 2020; Gruber & Ranganath, 2019 for reviews). This interpretation is supported by invasive animal research demonstrating a link between VTA-NAcc pathways and motivation-based learning and behaviour (Berridge, 2012; Vancraeyenest et al., 2020; Yang et al., 2018). Together, these findings support the view that intrinsic mesolimbic functional connectivity not only has a functional role in determining an individual’s motivation to seek and explore new information, but also consequently leads to better memory of particularly salient information. However, previous theoretical frameworks have suggested that dopaminergic projections from the VTA to the hippocampus are responsible for curiosity-related (Gruber & Ranganath, 2019) as well as reward- and novelty-driven memory enhancements (Düzel et al., 2010; Lisman & Grace, 2005; Shohamy & Adcock, 2010). Consistent with these theoretical ideas, fMRI studies have shown that task-based increases in VTA-hippocampal functional connectivity predict reward- and novelty-related memory enhancements (Adcock et al., 2006; Cowan et al., 2021; Gruber et al., 2016; Murty & Adcock, 2014) and curiosity-driven neuromodulation of the VTA on hippocampal states promotes subsequent memory formation (Poh et al., 2022). Based on these theoretical ideas and experimental findings, it might be considered surprising that no functional connections between VTA and hippocampus were associated with curiosity-driven memory enhancements in the present study. There are two explanations for why this relationship was not observed. First, VTA-hippocampal as compared to VTA-NAcc connections are more sparse and have a more precise innervation topography (Gasbarri et al., 1994; Zubair et al., 2021), making it more difficult to measure VTA-hippocampal functional connectivity with a seed-based resting-state fMRI analysis. Second, the present finding is in line with recent research suggesting that memory enhancements based on intrinsic curiosity and extrinsic reward share some commonalities but might rely on different functional and structural brain networks (Duan et al., 2020; Elliott et al., 2022; Meliss & Murayama, 2022). For instance and potentially consistent with our findings, Meliss & Murayama (2022) found that changes in VTA-hippocampal functional connectivity from pre-to post-learning rest periods were associated with externally rewarded but not curiosity-driven memory enhancements. Consequently, individual differences in intrinsic functional connectivity within the mesolimbic system, but not mesolimbic-hippocampal functional connectivity, might be the key determinant of individual differences in the magnitude of curiosity-driven memory enhancements. If this interpretation holds true in future studies, the PACE framework (Gruber & Ranganath, 2019), which currently suggests that the influence of the mesolimbic dopaminergic system on the hippocampus are necessary for the enhancement of curiosity-driven memory, needs to be adjusted.

The double dissociation between mesolimbic and cingulo-hippocampal functional connectivity respectively predicting curiosity-related and IPE-driven memory enhancements helps to disentangle the contrary predictions about the neural underpinnings of IPEs. The information-as-reward hypothesis suggests that IPEs – similar to reward prediction errors and curiosity – are dependent on reward circuitries, such as the mesolimbic system, indicating that information in itself can be rewarding (Marvin & Shohamy, 2016; van Lieshout et al., 2020). In contrast, the PACE framework pointed out the important role of the ACC and hippocampus in the detection and processing of prediction errors (Gruber & Ranganath, 2019). The findings of the present study are in line with the PACE framework showing that intrinsic cingulo-hippocampal but not mesolimbic functional connectivity predicted IPE-driven memory enhancements. Furthermore, they provide an addition to the PACE framework, which implies that the ACC and hippocampus are not only important for the detection and processing of prediction errors but the strength of their functional connection also leads to enhancement of memory performance related to prediction errors. Further evidence comes from a study showing that detection of prediction errors by the ACC triggers hippocampal activation, leading to memory replay and consolidation (Momennejad et al., 2018). Both findings are in line with theoretical frameworks pointing out the functional role of the ACC in signalling the degree to which information is unexpected or surprising, leading to adjustments in cognitive control that resolve the detected conflict (Alexander & Brown, 2011; Vassena et al., 2020), and in orchestrating attention and cognitive control processes in memory recall (Weible, 2013). Moreover, indirect projections from the ACC to the ventral hippocampus have been shown to be crucial for the generalisation of contextual memories in animals (Bian et al., 2019), making initially hippocampus-dependent memories increasingly dependent on cortical regions, such as the ACC, over time (see Frankland & Bontempi, 2005 for review). Consequently, the strength of cingulo-hippocampal functional connectivity might influence the successful integration of newly encountered, IPE-related information into existing neocortical memories (Audrain & McAndrews, 2022; Gruber et al., 2018; Takehara-Nishiuchi, 2020). In a study that used a comparable task design to the trivia paradigm, prediction errors were calculated as confidence in correct (positive prediction error) and incorrect answers (negative prediction error) to questions about previously studied information (Pine et al., 2018). In that study, both the valence and magnitude of prediction errors correlated positively with activation in the striatum, including the NAcc, and the cingulate cortex. Based on these findings and the discovered relationship of intrinsic cingulo-hippocampal functional connectivity and IPE-related memory enhancements in the present study, we would expect similar cingulo-hippocampal functional connections and neural activation in task-based assessments in the trivia paradigm. Furthermore, given that curiosity-related and IPE-driven memory enhancements were associated with distinct intrinsic functional connectivity, these findings suggest that memory formation based on curiosity and IPEs might be dependent on different brain networks within the cortico-mesolimbic dopaminergic circuit.

The association of distinct intrinsic functional connectivity with curiosity-driven and IPE-related memory enhancements indicates that neural fingerprints can indicate the degree to which individuals benefit from one or the other incentive salience, respectively, adding to previous research predicting behavioural performance from resting-state neuroimaging data (Baldassarre et al., 2012; Gerraty et al., 2014). Moreover, these findings aid the idea that intrinsic functional networks measured at rest support task-related cognitive processes (Cole et al., 2014; Gratton et al., 2018). Similarly to the separable memory-enhancing effects of reward and strategic encoding (Cohen et al., 2019), curiosity and IPEs represent distinct cognitive mechanisms that depend on dissociable intrinsic neural networks but can enhance memory within the same paradigm. Recently, a study that used a whole-brain analysis approach failed to show a relationship between intrinsic functional connectivity and emotion-driven memory enhancements (Kandaleft et al., 2022). Regarding the double dissociation of the present study, it is possible that memory enhancements based on curiosity and IPEs are more robustly influenced by intrinsic functional connectivity. Alternatively, the theory-driven selection of ROIs for functional connectivity analysis as it was used in the present study might be a helpful approach to derive meaningful brain-behaviour associations. In line with the finding that intrinsic functional connections between the PFC and hippocampus predict an individual’s learning behaviour (Gerraty et al., 2014), the present study demonstrated a relationship between intrinsic LPFC-hippocampal functional connectivity and overall memory performance. This finding is consistent with ample research demonstrating higher encoding-related PFC and hippocampus activations that are interpreted to reflect memory control processes in the PFC that are necessary to support hippocampus-dependent memory formation (see Anderson et al., 2016; Kim, 2011; Ranganath, 2010; Simons & Spiers, 2003 for reviews).

In conclusion, the present study provides evidence for dissociable intrinsic functional connections that determine inter-individual differences in memory enhancements, thereby informing theoretical frameworks on curiosity and curiosity-driven memory, but also highlighting the need to account for inter-individual differences in how curiosity and prediction errors affect learning in applied settings.

## ACKNOWLEDGEMENTS

We thank Alison Cooper, Vera Dehmelt, John Evans, Peter Hobden, and Duarte Pereira for help with MRI scanning and data collection. This work was supported by a Wellcome Trust ISSF Consolidator Award (AC1710IF09) and a German Research Foundation Research Fellowship (442588275) awarded to K.C.J.E.; a PhD studentship funded by the Cardiff University School of Psychology awarded to A.V.; a Wellcome Trust Strategic Award (104943/Z/14/Z) granted to K.S.G.; a BBSRC grant (BB/V008242/1) awarded to A.D.L.; and a Wellcome Trust and Royal Society Sir Henry Dale Fellowship (211201/Z/18/Z) awarded to M.J.G. For the purpose of Open Access, the authors have applied a CC-BY public copyright licence to any Author Accepted Manuscript version arising from this submission.

## AUTHOR CONTRIBUTIONS

K.C.J.E., A.V., and M.J.G. designed research; K.C.J.E., and A.V. performed research, K.C.J.E. analysed the data; K.S.G, A.D.L., and M.J.G. supervised research; K.C.J.E. drafted the manuscript; K.C.J.E., K.S.G, A.D.L., and M.J.G. wrote the paper.

